# Mechanistic Links Between Metabolic Dysfunction-Associated Steatotic Liver Disease and Heart Failure with Preserved Ejection Fraction in a Mouse Model

**DOI:** 10.1101/2025.11.02.686066

**Authors:** Niveen Rweished, Yair Rokach, Elchanan Parnassa, Suzan Muhamad Abedat, Aseel Bsoul, Dean Nachman, Offer Amir, Rifaat Safadi, Rabea Asleh

## Abstract

**Background:** Metabolic dysfunction-associated steatotic liver disease (MASLD) commonly coexists with heart failure with preserved ejection fraction (HFpEF), yet the mechanisms linking hepatic steatosis to diastolic dysfunction remain unclear.

**Methods:** A combined murine model of pressure overload using transverse aortic constriction (TAC) and diet-induced MASLD was developed to investigate the liver-heart interaction in HFpEF. Cardiac function and structure were assessed by echocardiography and histopathology. Hepatic transcriptomics and cardiac metabolomics were integrated to identify molecular pathways underlying cardiac remodeling and diastolic dysfunction.

**Results:** While left ventricular ejection fraction remained preserved, HFpEF-MASLD mice exhibited significantly worse diastolic function (lower septal è velocity and higher E/è ratio), greater left ventricular hypertrophy, and more extensive myocardial fibrosis compared with HFpEF alone. Immunofluorescence demonstrated augmented myocardial inflammation with increased CD3^+^ T-cell and CD68^+^ macrophage infiltration in the combined HFpEF-MASLD group. Transcriptomic analysis demonstrated marked down-regulation of genes implicated in retinoic acid signaling, confirmed by reduced expression of retinoic acid receptors (RARα, RARβ) and retinol dehydrogenases (RDH) in both hepatic and cardiac tissues. Cardiac metabolomics revealed suppression of arginine biosynthesis, the obligate substrate for endothelial nitric oxide (NO) synthase, suggesting a potential link to reduced NO-mediated vascular and myocardial signaling.

**Conclusions:** MASLD aggravates HFpEF through converging inflammatory and metabolic derangements. Disruption of retinoic acid and arginine-NO pathways may represent an important mechanistic link between hepatic steatosis and diastolic dysfunction that warrants further mechanistic and translational investigation.

## Introduction

Metabolic dysfunction-associated steatotic liver disease (MASLD) is now recognized as the most prevalent cause of chronic liver disease globally, which may ultimately cause severe liver-related complications, including cirrhosis and hepatocellular carcinoma (1). Importantly, cardiovascular disease represents the leading cause of mortality in MASLD patients (2). Affecting approximately 38% of the general population, the global health burden of MASLD is escalating rapidly, largely driven by the increasing prevalence of obesity and type 2 diabetes mellitus (T2DM) (1,3). These metabolic comorbidities not only heighten the risk of MASLD development but also accelerate its progression to more advanced liver pathologies (1).

In recent years, significant attention has shifted towards the emerging link between MASLD and cardiovascular diseases, including coronary artery disease and heart failure (HF), among others (4–7). Numerous epidemiological studies have consistently identified MASLD as an independent and significant risk factor for HF development (6–9). Notably, a study by Chang and colleagues reported that 1.6% of a large MASLD patient cohort progressed to HF, with a striking 76% of these cases presenting as heart failure with preserved ejection fraction (HFpEF) (8). This distinctive interplay between the heart and the liver is frequently referred to as “cardio-hepatic interactions” (9).

HFpEF affects nearly half of all patients with HF worldwide, with its prevalence continuously rising (10). HFpEF is increasingly recognized as a multi-system syndrome influenced by chronic comorbidities such as obesity, hypertension, and T2DM, highlighting its complex pathophysiology and extracardiac involvement (11). A critical component in HFpEF development and deterioration is dysregulated immune response and chronic inflammation, both locally within the myocardium and systemically (12–14). In obese HFpEF patients, stronger correlations are observed between body weight, cardiac filling pressures, ventricular remodeling, and adverse hemodynamics (15,16). Weight gain and excess central adiposity further contribute to ventricular stiffening and myocardial dysfunction (15,17). Furthermore, systemic inflammation and adipogenesis, prominent features in liver disorders like MASLD and metabolic dysfunction-associated steatohepatitis (MASH), are increasingly associated with adverse cardiac effects and HFpEF progression (18–20). These observations suggest a causative relationship between liver health and HF, potentially driving specific HFpEF phenotypes (7,9).

Interestingly, recent conceptual frameworks propose that adipose tissue-derived signaling may represent a unifying mechanism linking obesity, MASLD and HFpEF. The adipokine hypothesis postulates that expansion of visceral and epicardial adipose tissue alters the balance of secreted factors toward proinflammatory and profibrotic adipokines (e.g., leptin, resistin, TNF-α) while suppressing protective adipokines (e.g., adiponectin, apelin). This imbalance promotes systemic inflammation, endothelial dysfunction, myocardial fibrosis, and diastolic impairment (21). Given the shared metabolic and inflammatory milieu, MASLD may function as a hepatic extension of this adipose-cardiac axis, releasing hepatokines and inflammatory mediators that amplify myocardial stress, oxidative injury, and fibrosis. This emerging paradigm strengthens the concept of a causal continuum between steatotic liver disease and HFpEF progression.

Building on these insights, growing evidence points to MASLD as a direct driver of specific HFpEF phenotypes, which are thought to represent a continuum from mild to severe disease (7). These proposed MASLD-driven HFpEF phenotypes include: 1) obstructive HFpEF, primarily linked to preload reserve failure; 2) metabolic HFpEF, characterized by metabolic syndrome with inflammation, endothelial dysfunction, and excess pericardial adipose tissue; and 3) advanced liver disease/cirrhosis HFpEF, associated with increased cardiac output due to portosystemic and arteriovenous shunts (22). Among these, a prominent metabolic phenotype of HFpEF has been identified through phenomapping studies, characterized by a high prevalence of obesity, chronic kidney disease, left ventricular hypertrophy, and significant cardiac and pulmonary vascular dysfunction (23,24). This phenotype exhibits features directly associated with MASLD, such as elevated proinflammatory biomarkers and liver fibrosis, strongly supporting a pathophysiological continuum between MASLD and HFpEF (23,24). This continuum is likely mediated by multiple shared mechanisms, including systemic inflammation, endothelial dysfunction, and perturbed systemic metabolism (19–22,25).

While the epidemiological connection between MASLD and HF, particularly HFpEF, is well-established, the precise molecular and cellular mechanisms underpinning this critical inter-organ crosstalk remain largely unknown. Identifying these specific mechanistic links is crucial for understanding disease pathogenesis and for developing targeted therapeutic strategies for patients suffering from both MASLD and HFpEF. Given these observations, we hypothesized that specific mechanistic pathways mediate this crucial communication between the liver and the heart, thereby contributing to HFpEF progression in the setting of MASLD.

## Materials and Methods

### Experimental design

Male C57BL/6 mice, aged 9 to 12 weeks, were obtained for this study. All animal experiments were approved by the local Ethics Committee and conducted in strict accordance with the guidelines outlined in “The Guide for the Care and Use of Laboratory Animals, published by the US National Institutes of Health (NIH Publication No. 85-23).” Mice were housed under controlled conditions with a 12-hour light/12-hour dark cycle (6:00 AM to 6:00 PM).

### Study protocol and experimental groups

The experimental timeline for inducing HFpEF and MASLD is depicted in ***Fig. 1A***. Sixty mice were randomly assigned to one of four experimental groups: (1) Sham group (n=10): Mice were maintained on a standard chow diet (2918, Teklad) throughout the study and underwent a sham surgical procedure (thoracotomy without aortic constriction) at the beginning of the experiment; (2) MASLD group (n=10): Mice were fed with a high-fat diet (HFD) containing approximately 60% of kilocalories from fat (TD.06414, Envigo) for 12 weeks. They also underwent a sham surgical procedure at the beginning of the experiment; (3) HFpEF group (n=20): Mice were maintained on a standard chow diet and underwent transverse aortic contraction (TAC) surgery at the beginning of the experiment; and (4) combined HFpEF and MASLD mice (HFpEF-MASLD group) (n=20): Mice were fed with HFD for 12 weeks and underwent TAC surgery at the beginning of the experiment.

**Figure 1.**
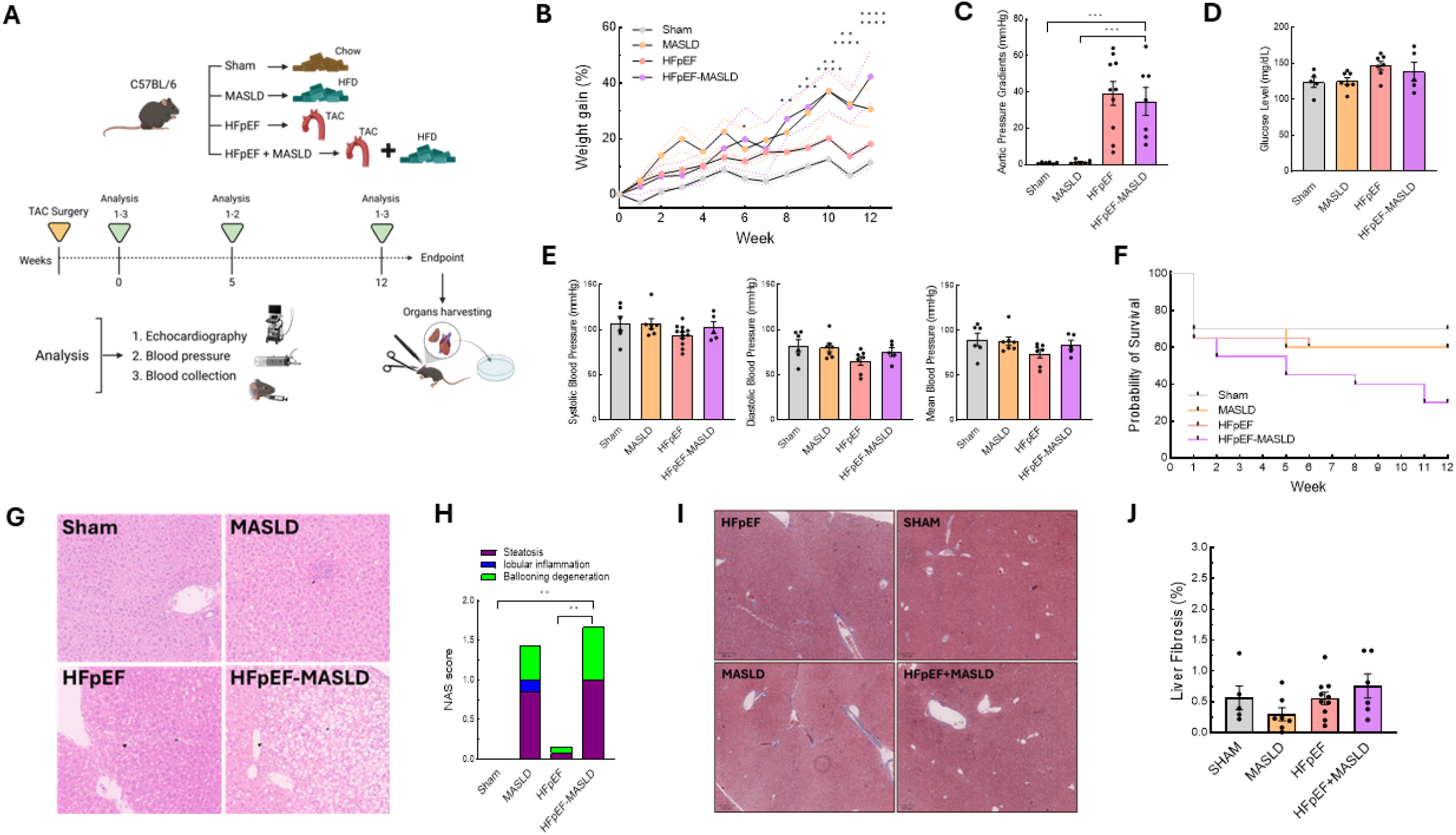
Experimental groups during the period of 12 weeks. **A**, Experimental design. **B**, Weight gain. Body weight was measured on the same day of every week for 12 weeks. *n* = 6-13 mice per group. (Mixed-effects analysis followed by Dunnett’s multiple comparisons test. *P <0.05 HFpEF-MASLD versus Sham; 6 weeks, **P <0.01 HFpEF-MASLD versus Sham; 8 weeks, ***P <0.001 HFpEF-MASLD versus Sham, *P <0.05 HFpEF-MASLD versus HFpEF; 9 weeks, ****P <0.0001 HFpEF-MASLD versus Sham, **P <0.01 HFpEF-MASLD versus HFpEF; 10 weeks, ****P <0.0001 HFpEF-MASLD versus Sham, **P < 0.01 HFpEF-MASLD versus HFpEF; 11 weeks, ****P <0.0001 HFpEF-MASLD versus Sham, ****P <0.0001 HFpEF-MASLD versus HFpEF; 12 weeks). **C**, Mice aortic pressure gradients (mmHg) were measured by echocardiography at week 5. n = 7-10 per group. **D**, Blood glucose level was measured at week 12. n = 5-7 per group. **E**, Systolic, diastolic, and mean blood pressure (mmHg) was measured at week 12. *n* = 5-10 mice per group. **F**, Kaplan-Meier survival curves over a 12-week period for SHAM, MASLD, HFpEF, and HFpEF-MASLD groups. Log-rank test for trend was significant (P <0.01). *n* = 7-10 mice per group. **G**, Representative light microscope images of liver tissue after H&E staining. Magnification x20. Sham and HFpEF mice showed no hepatic steatosis or ballooning degeneration, while MASLD and HFpEF-MASLD mice showed significant hepatic steatosis (arrows) and ballooning degeneration (arrowheads). **H**, Quantification of hepatic histopathology using the NAFLD Activity Score (NAS), which includes steatosis (purple), lobular inflammation (blue), and ballooning degeneration (green). NAS score was significantly higher in the MASLD and HFpEF-MASLD mice compared to the other groups (**P <0.001 by 2-way ANOVA and Dunnett’s multiple comparisons corrections analysis). **I**, Representative light microscope images of liver tissue after Masson’s trichome staining. The scale bar is 20 µm. Magnification x6. **J**, Fibrosis per HPF. *n =* 4-9 mice per group. The results are expressed as the mean ±SEM.

HFpEF was surgically induced by TAC. The TAC surgical procedure was performed based on previously established protocols with minor modifications (26,27). All surgical interventions (TAC or sham) were performed during the initial week of the experiment. Investigators were blinded to the treatment assignment of individual animals during the experiments and outcome assessments.

### Echocardiography and Doppler imaging

Transthoracic echocardiography was performed to assess cardiac structure and function using a VisualSonics Vevo 3100 system equipped with an MX550D transducer. Anesthesia was induced with 5% isoflurane and confirmed by the lack of response to firm pressure on a hindpaw. During echocardiogram acquisition, mice were placed on a temperature-controlled platform to maintain body temperature, and isoflurane was reduced to 1.0-1.5% and adjusted to maintain a stable heart rate between 400-500 beats per minute. The effectiveness of the TAC procedure was confirmed by measuring the transaortic pressure gradient using continuous-wave Doppler echocardiography at the level of the aortic constriction.

Left ventricular systolic function was obtained from short-axis M-mode scans acquired at the midventricular level, identified by the presence of papillary muscles. Parameters measured included left ventricular end-diastolic diameter (LVEDD), left ventricular end-systolic diameter (LVESD), left ventricular mass, and left ventricular ejection fraction (LVEF). Diastolic function was assessed from apical four-chamber views using pulsed-wave (PW) and tissue Doppler imaging (TDI) at the mitral valve level. Key diastolic parameters collected included: peak early (E) and late (A) diastolic Doppler blood inflow velocities across the mitral valve (E/A ratio), peak early diastolic tissue Doppler myocardial relaxation velocity at the mitral valve annulus (è) and early filling deceleration time.

Global longitudinal strain (GLS) was calculated from B-mode traces acquired from the parasternal long-axis view using Vevo Strain software (Visual Sonics) with a speckle-tracking algorithm. Longitudinal strain values are inherently negative, reflecting myocardial fiber shortening. Images were selected based on high frame rates and clear visualization of both endocardial and epicardial left ventricular wall borders. Borders were manually traced, and semi-automated strain analysis was performed. Average peak global strain values were obtained from six independent anatomical segments of the left ventricle.

### Non-invasive blood pressure measurement

Systolic and diastolic blood pressure were measured non-invasively in awake mice using a tail-cuff technique with a CODA instrument (Kent Scientific). Mice were habituated to individual holders on a temperature-controlled platform (32 °C). Recordings were performed under steady-state conditions, ensuring stable measurements for analysis.

### Histology and immunofluorescence staining

At the study endpoints, mouse cardiac ventricles and liver tissues were collected, fixed in 4% paraformaldehyde for 48 hours, and processed for routine paraffin embedding. Sections were cut at 5-μm thickness. For histological staining, representative sections were stained with Hematoxylin and Eosin (H&E) for general morphology and Masson’s Trichrome for collagen deposition and fibrosis. For immunofluorescence staining, paraffin-embedded cardiac ventricle sections were deparaffinized and subjected to antigen retrieval using hot citrate buffer (Antigen Unmasking Solution, Vector Laboratories). Non-specific binding was blocked with 3% bovine serum albumin (BSA) in phosphate buffered saline (PBS). Sections were then incubated overnight at 4°C with the following primary antibodies: Rat anti-CD3 monoclonal antibody (1:100; Abcam, ab11089) and rabbit anti-CD68 monoclonal antibody (1:100; Abcam, ab283654). After thorough washing, sections were incubated for 1 hour at room temperature with the following secondary antibodies diluted in CAS-Block (Jackson Immunoresearch Laboratories): Cy5-conjugated donkey anti-Rat IgG (1:100; Jackson Immunoresearch Laboratories) and Cy5-conjugated goat anti-Rabbit IgG (1:100; Jackson Immunoresearch Laboratories). Sections were then mounted using an anti-fade reagent supplemented with DAPI (4’,6-diaminide-2-phenylindole) for nuclear counterstaining (e.g., Cat# 17984-24, Biotrend or specific vendor for DAPI if part of the mounting medium kit). Images were acquired using Nikon fluorescent microscopy.

### RNA sequencing

Total RNA was extracted from the liver tissue of all experimental groups using The SMARTer Stranded Total RNA Sample Prep Kit - HI Mammalian (Takara Bio USA, Inc., catalog no. 634873). This kit facilitates the generation of indexed cDNA libraries suitable for next-generation sequencing (NGS) on Illumina platforms. The library preparation protocol was performed according to the manufacturer’s instructions, involving ribosomal RNA (rRNA) removal, first-strand cDNA synthesis, purification of first-strand cDNA using SPRI AMPure Beads, RNA-Seq library amplification by PCR, and final purification of the RNA-Seq library using SPRI AMPure Beads. RNA input ranging from 100 ng to 1 μg of total mammalian RNA was used. Reverse transcription involved incubation in a thermal cycler at 37°C for 30 min. Library validation and quality control were performed using the Agilent 2100 Bioanalyzer (Agilent Technologies). Libraries were pooled and sequenced on an Illumina NovaSeq 6000 platform to generate 122 bp single-end reads.

### Bioinformatic analysis of RNA-sequencing data

Raw sequencing reads were subjected to quality control using FastQC (version 0.11.9). Adapter removal and trimming of low-quality bases were performed with Trimmomatic (version 0.39). Cleaned reads were then aligned to the mouse reference genome GRCm39 (Ensembl release 106) using TopHat2 (version 2.1.1). Gene expression quantification was performed using HTSeq-count (version 2.0.1). Genes with fewer than 10 total reads across all samples were excluded. Normalization and differential gene expression analysis were conducted with DESeq2 (version 1.36.0) (28) in R (version 4.2.2). A negative binomial generalized linear model was fitted to the count data to identify genes significantly altered between experimental groups. Genes with an adjusted p-value < 0.05 (Benjamini-Hochberg corrected for multiple comparisons) were considered significantly differentially expressed.

Volcano plots were generated to visualize differentially expressed genes between specific groups (e.g., HFpEF vs. HFpEF-MASLD), with significantly upregulated and downregulated genes indicated by distinct colors. Hierarchical clustering and heatmaps were used to display the expression profiles of all significantly differentially expressed genes across relevant samples (R packages ggplot2 version 3.4.0 and heatmap version 1.0.12). Venn diagrams were constructed to show unique and shared significantly differentially expressed gene sets across selected pairwise group comparisons (Venny version 2.0.2).

For functional annotation, a protein-protein interaction (PPI) network of unique differentially expressed genes was constructed and visualized using the STRING database (version 11.5) with node colors indicating cluster identities. Additionally, functional annotation analysis was performed using the DAVID web tool (2021 update) (29) to identify enriched Gene Ontology (GO) terms and Kyoto Encyclopedia of Genes and Genomes (KEGG) pathways. Bar graphs illustrating the top enriched and significant biological terms, along with tables listing representative genes associated with each term, were generated.

### Metabolomics

Ventricular tissues from HFpEF and HFpEF-MASLD mice were collected at the study endpoint for metabolic profiling. The samples were homogenized using a Precellys 24 homogenizer with a cryolys temperature controller (Bertin Instruments) maintained at approximately 0°C (3 cycles of 30 seconds, with 30-second intervals between cycles). Homogenates were then centrifuged in the homogenization tubes at 13,000 RPM for 20 minutes at 4°C. The supernatant was collected into HPLC vails. For quality control, an aliquot from each sample was pooled to create a single Quality (QC) sample.

LC-MS metabolomics analysis was performed using a Dionex Ultimate 3000 high-performance liquid chromatography (HPLC) system coupled to an Orbitrap Q-Exactive Plus Mass Spectrometer (Thermo Fisher Scientific). The mass spectrometer was operated with a resolution of 70,000 at 200 m/z, utilizing electrospray ionization in the HESI source. Data acquisition was performed in polarity switching mode to enable detection of both positive and negative ions across a mass range of 70 to 1000 m/z. Mass accuracy for all detected metabolites was maintained below 5 ppm. Data acquisition was controlled by Thermo Xcalibur software.

Chromatographic separation was achieved using a ZIC-pHILIC column (SeQuant; 150 mm × 2.1 mm, 5 μm; Merck) equipped with a Sure-Guard filter (SS frit 0.5 μm). Five microliters (5 µL) of the tissue extracts were injected. Compounds were separated with a 15-minute mobile phase gradient. The gradient started at 20% aqueous phase (20 mM ammonium carbonate adjusted to pH 9.2 with 0.1% of 25% ammonium hydroxide) and 80% organic phase (acetonitrile). The organic phase was reduced to 20% acetonitrile over 15 minutes, followed by a re-equilibration phase. The flow rate was maintained at 0.2 mL/min, and the column temperature at 45°C, for a total run time of 26 min per sample.

Raw LC-MS data files were processed using Compound Discoverer 3.3.2.31 software (Thermo Fisher Scientific). This included peak picking, alignment, and putative metabolite annotation using an in-house library (30). For metabolomic profiling and pathway analysis, processed data from heart tissue of HFpEF and HFpEF-MASLD mice were subjected to various statistical and visualization methods. These included Partial Least Squares-Discriminant Analysis (PLS-DA) to identify group separation, Volcano plots to visualize differentially abundant metabolites between groups, and Heat Maps to display metabolite expression patterns across samples. Pathway analysis was performed using the MetaboAnalyst web tool (version 6.0) to identify significantly enriched metabolic pathways (31).

### Statistical Analysis

All data are presented as mean ± standard error of the mean (SEM) unless otherwise specified. For comparisons between two groups, Student’s t-tests were used for normally distributed data and Mann-Whitney U tests for non-normally distributed data. Differences among three or more groups were analyzed using one-way analysis of variance (ANOVA) followed by Tukey’s post-hoc test for multiple pairwise comparisons of parametric data, while Kruskal-Wallis tests followed by Dunn’s multiple comparison test were employed for non-parametric data. For repeated measures, two-way ANOVA with Dunnett’s post-hoc test was used to compare groups over time. Survival curves were compared by log-rank test. For RNA-sequencing data, differential gene expression analysis was performed as described in the “Bioinformatic analysis of RNA sequencing data” section, using a negative binominal generalized linear model implemented in DESeq2 (28). Statistical significance for gene expression was determined using an adjusted p-value <0.05 after Benjamini-Hochberg correction for multiple comparisons. Metabolomics data underwent PLS-DA for multivariate analysis and identification of differentially abundant metabolites. Pathway analysis was conducted using the MetaboAnalyst web tool (version 6.0) (31), as detailed above. For comparisons, a two-tailed p-value <0.05 was considered statistically significant. Statistical analyses were conducted using GraphPad Prism software (version 10.0) and R (version 4.2.2; R Core Team, 2022). Experimental mice were randomly assigned to each experimental or control group. Data acquisition and analysis were performed by investigators blinded to the experimental group assignments.

## Results

### Systemic phenotype of mouse models

During the 12-week experimental period, the distinct systemic phenotypes of each experimental group were characterized. As anticipated, both MASLD and HFpEF-MASLD mice developed significant obesity, evidenced by increased body weight compared to the Sham and HFpEF groups (***Fig. 1B***). The successful induction of TAC in the HFpEF and HFpEF-MASLD groups was confirmed by a significant increase in the transaortic pressure gradient, as measured by echocardiography, hence resulting in an increased left ventricular afterload, as compared to sham-operated controls (***Fig. 1C***). Plasma glucose levels were similar among the study groups (***Fig. 1D***). Importantly, non-invasive tail-cuff measurements of blood pressure revealed no significant changes in either systolic or diastolic blood pressure across any experimental group at baseline or throughout the study duration, resulting in similar blood pressure measurements among all groups (***Fig. 1E***). Survival rates at 12 weeks post-surgery were significantly decreased in HFpEF-MASLD mice (30% survival), while the sham group had 70% survival and both MASLD and HFpEF groups had 60% survival (p<0.01) (***Fig. 1F***).

MASLD was histologically confirmed in both the MASLD and HFpEF-MASLD groups using non-alcoholic fatty liver disease (NAFLD) activity score (NAS) (***Fig. 1G, H***). Both MASLD-only and HFpEF-MASLD mice consistently exhibited mild steatosis (NAS score ≤2), lobular inflammation, and ballooning degeneration. In contrast, the chow-diet groups (Sham and HFpEF) showed no accumulation of hepatic fat, inflammation, or ballooning degeneration (***Fig. 1G, H***). Liver fibrosis, a key marker for advanced liver damage, was assessed by Masson’s Trichrome staining, showing no significant fibrosis in the MASLD model and without differences among the experimental groups (***Fig. 1I, J***). These results indicate that the MASLD model generated in this study represents early stages of the disease that typically involve hepatic steatosis and balloon degeneration but without significant fibrosis.

### MASLD exacerbates diastolic dysfunction in HFpEF

To evaluate cardiac function, high-frequency echocardiography was performed on anesthetized mice, with heart rates maintained in the range of 400-500 beats per minute (***Fig. 2A***). Longitudinal echocardiography revealed preserved systolic function across all treatment groups, as evidenced by normal left ventricular ejection fraction (LVEF) (***Fig. 2B***). Diastolic function, however, showed significant impairment in the combined HFpEF and MASLD model compared to all other groups. Specifically, early diastolic mitral annular velocity (e′), obtained by tissue Doppler imaging and reflecting myocardial relaxation (***Fig. 2C***), was significantly reduced in the HFpEF-MASLD group. Correspondingly, the ratio of the peak early mitral inflow velocity (E) over è (E/è), which reflects left ventricular end-diastolic filling pressure (***Fig. 2D***), was significantly elevated in the HFpEF-MASLD group. Additionally, the E/A ratio was higher in the HFpEF-MASLD group compared to the HFpEF-alone group (***Fig. 2E***).

**Figure 2.**
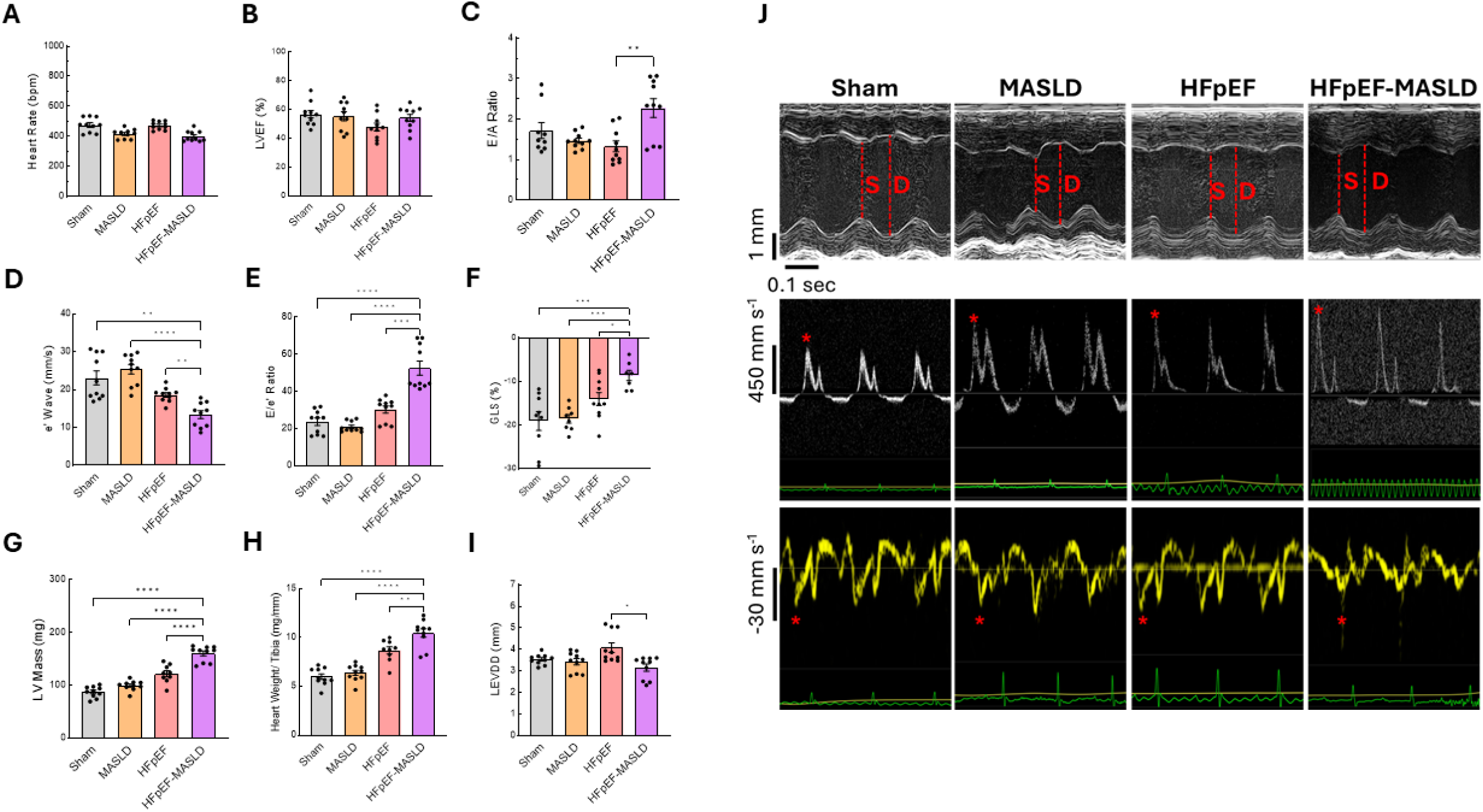
Effects of MASLD on systolic and diastolic function in HFpEF mice. **A**, Heart rate (bpm). **B**, Left ventricular ejection fraction (LVEF). **C**, Ratio between early mitral (E) and late (A) ventricular filling velocities. **D**, Septal mitral annulus velocity by tissue Doppler (e’). **E**, Ratio between mitral inflow velocity (E) and e′ (E/e′). **F**, Left ventricular global longitudinal strain (GLS). **G**, Left ventricular mass assessed by echocardiography. **H**, Left ventricular weight normalized to tibia length (mg/mm). **I**, Left ventricular internal diameter measured by echocardiography. **J**, Representative tracings: M-mode (top; S, systole; D, diastole), pulsed-wave Doppler at the mitral inflow (middle; asterisk denotes E-wave), and tissue Doppler at the septal annulus (bottom; asterisk denotes e′). ECG trace (green) and respiratory trace (yellow) shown below. Bars represent mean ± SEM with individual mice as dots. n=7-10 mice per group for panels A-I. Statistical significance was tested by one-way ANOVA or Kruskal-Wallis with appropriate multiple-comparison corrections tests as described in Methods.

Left ventricular GLS, reflecting regional myocardial contraction in the long-axis direction, was significantly reduced in the HFpEF-MASLD group as compared to the control groups (***Fig. 2F***), indicating impaired myocardial mechanics. Assessment of cardiac hypertrophy revealed that ventricular weight was significantly greater in both the HFpEF and HFpEF-MASLD groups compared to the sham and MASLD groups, indicating a hypertrophic phenotype in these TAC-challenged models. More prominent LV hypertrophy was observed in HFpEF-MASLD mice compared to the other groups (***Fig. 2G***). Similarly, the heart weight normalized to tibia length was significantly increased in the HFpEF-MASLD group compared to the other groups (***Fig. 2H***). Consistent with the marked LV hypertrophy seen in the HFpEF-MASLD group, these mice also exhibited a significantly more decreased LVEDD as compared to HFpEF-alone group (***Fig. 2I***). Representative M-mode (***Fig. 2J*, *top***), pulse-wave Doppler (***Fig. 2J*, *middle***), and tissue Doppler (***Fig. 2J*, *bottom***) tracings illustrate key echocardiographic parameters for structural and diastolic function assessment, highlighting differences among the study groups.

### MASLD exacerbates myocardial fibrosis and inflammation in HFpEF

Cardiac fibrosis, characterized by excessive collagen deposition in the extracellular matrix, is a key component of impaired cardiac muscle relaxation and directly contributes to HFpEF pathophysiology (32,33). Histological analysis using Masson’s Trichrome staining revealed a significant increase in fibrotic areas within the myocardium of the combined HFpEF and MASLD model compared to both the sham and MASLD groups (***Fig. 3A, B***). Given the elevated levels of myocardial fibrosis observed in the HFpEF-MASLD group, we next investigated differences in inflammatory cell infiltration in the myocardium across the study groups. Immunofluorescence staining demonstrated a significant increase in CD3-positive cells (lymphocytes) (***Fig. 3C, D***) and CD68-positive cells (macrophages) (***Fig. 3E, F***) in the HFpEF-MASLD group as compared to the sham, MASLD, and HFpEF groups. These results suggest more pronounced inflammation within the myocardium of HFpEF-MASLD mice compared with HFpEF-alone mice.

**Figure 3.**
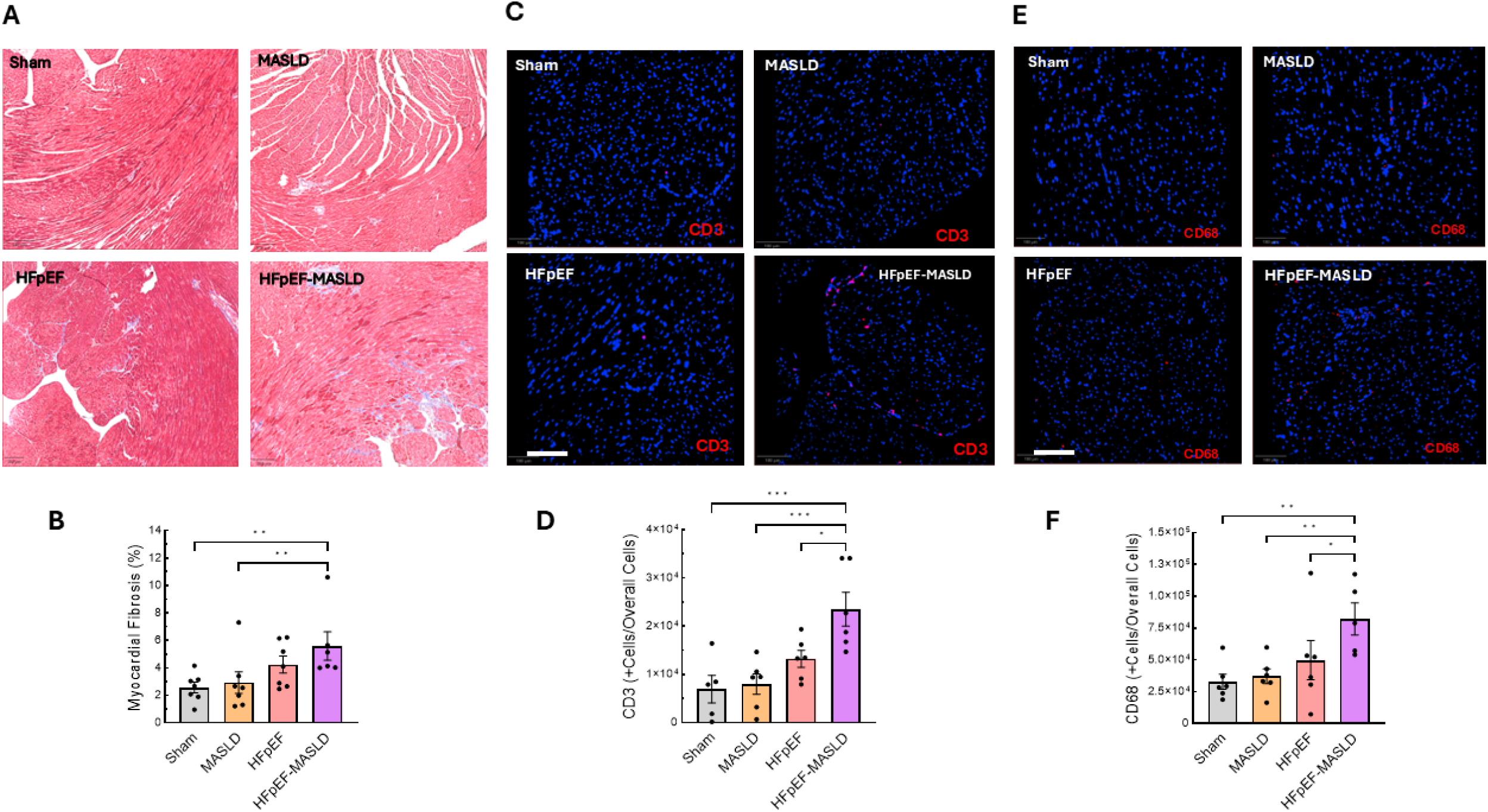
Light microscopy and immunofluorescence microscopy of myocardial tissues of the different experimental groups. **A**, Representative light microscope images of left ventricular tissue after Masson’s trichome staining. n = 6-7 mice per group. Magnification x14. **B**, Fractional fibrosis area (% of left ventricular tissue). n = 6-7 mice per group. **C**, Representative immunofluorescence staining for CD3 (T cells, red) with nuclear counterstain DAPI (blue). *n* = 6 mice per group. Magnification x20. Scale bar, 100 µm. **D**, Quantification of CD3^+^ cells per tissue area. *n* = 6 mice per group. **E**, Representative immunofluorescence for CD68 (macrophages, red) with nuclear counterstain DAPI (blue). *n* = 6 mice per group. Magnification x20. Scale bar, 100 µm. **F**, Quantification of CD68^+^ cells per tissue area. *n* = 6 mice per group. The results are expressed as the mean ±SEM.

### Hepatic transcriptomic profiling reveals retinoid and inflammatory pathways associated with HFpEF progression in HFpEF-MASLD

To elucidate hepatic mechanisms contributing to HFpEF progression in the setting of MASLD, we performed comprehensive RNA-sequencing-based transcriptomic profiling of liver tissue, comparing HFpEF-MASLD to the HFpEF-alone groups. A volcano plot analysis identified a total of 58 differentially expressed genes between the two groups (FDR <0.05), of which 26 were significantly upregulated and 32 significantly downregulated in the HFpEF-MASLD group relative to the HFpEF-alone group (***Fig. 4A***). The overall expression profiles of these significantly altered genes are further visualized in a heatmap, illustrating distinct clustering of samples according to disease group (***Fig. 4B***). To identify unique genes specifically associated with the combined HFpEF-MASLD pathology in the liver, Venn-diagram analysis was conducted across all pairwise group comparisons (MASLD vs sham, HFpEF vs sham, HFpEF-MASLD vs HFpEF). This analysis identified 40 genes specifically dysregulated only in the HFpEF-MASLD group, representing a liver transcriptomic signature exclusive to the combined phenotype (***Fig. 4C***).

**Figure 4.**
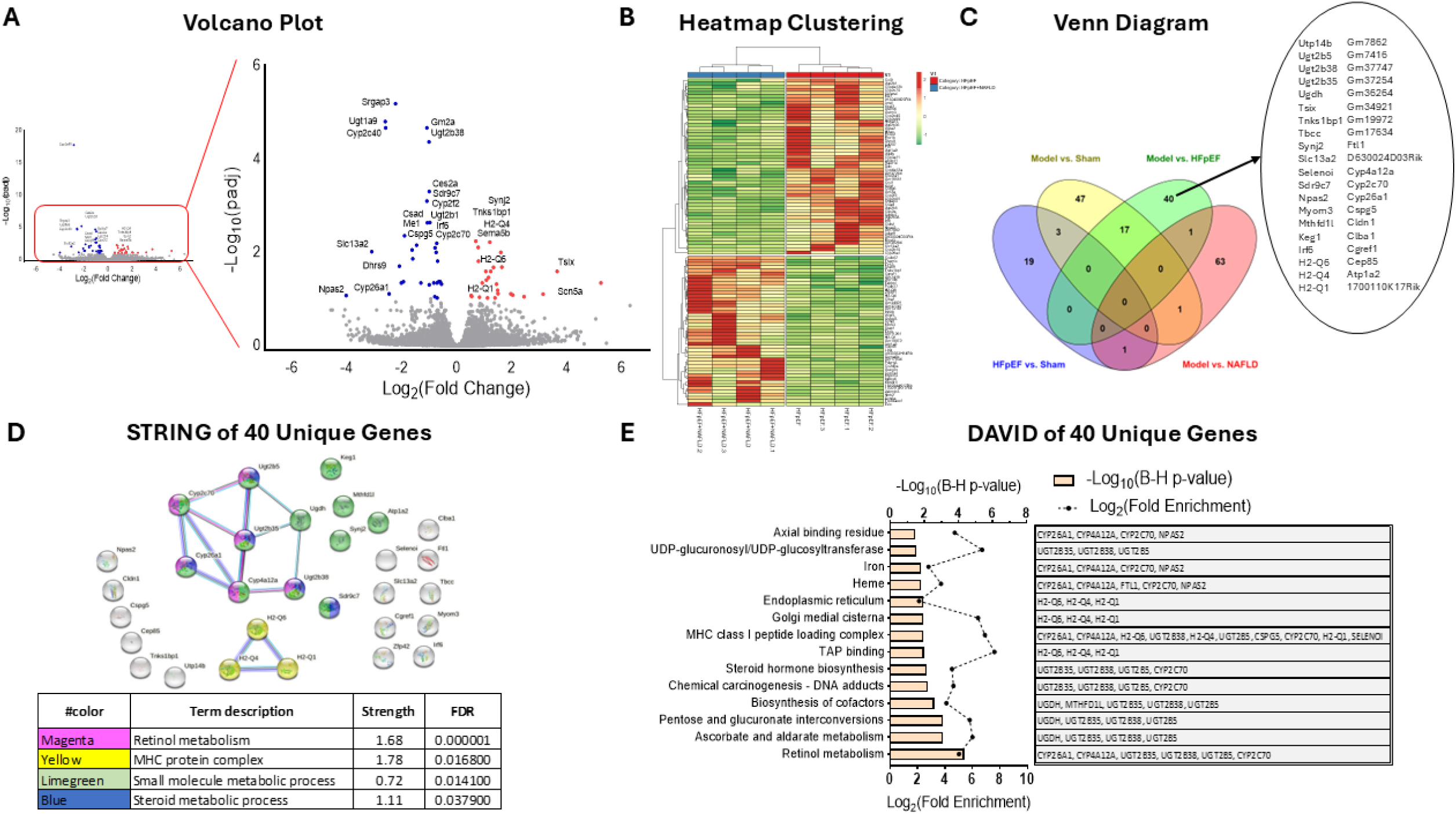
Transcriptomic profiling and functional enrichment analysis revealed distinct hepatic gene expression signatures in HFpEF-MASLD vs. HFpEF mice. **A**, Volcano plot of differentially expressed genes (DESeq2). X-axis: log_2_(fold change) for HFpEF-MASLD vs. HFpEF; Y-axis: −log_10_(adjusted p). Red/blue dots denote significantly up-/down-regulated genes (FDR <0.05, Benjamini-Hochberg). **B**, Heatmap of all significantly differentially expressed genes between HFpEF and HFpEF-MASLD groups. Each column represents an individual sample; rows represent genes; values are row-wise z-scores; hierarchical clustering as described in Methods. **C**, Venn diagram displaying the number of unique and shared significantly differentially expressed genes across selected pairwise group comparisons (HFpEF vs Sham, HFpEF-MASLD vs Sham, HFpEF vs MASLD, HFpEF-MASLD vs MASLD). **D**, Protein-protein interaction (PPI) network of 40 unique differentially expressed genes generated with STRING v11.5 (default settings) (top). Node colors indicate cluster identity listed in the enrichment score table (bottom). Line thickness reflects interaction confidence. The table lists cluster-level enrichment terms. **E**, Functional annotation analysis of the 40 unique differentially expressed genes using the DAVID database (2021 update). Bar graph shows the top enriched and significant biological GO/KEGG terms (multiple-testing corrected) (left), and the adjacent table lists representative genes associated with each term (right).

To assess the functional relationship and potential clustering among these 40 unique genes, a protein-protein interaction (PPI) network analysis was constructed using the STRING database. This analysis revealed significant functional clustering of certain gene sets (p <0.001 by STRING enrichment; ***Fig. 4D*, *top)***. Two major interaction modules were identified: one enriched for genes related to lipid and retinoid metabolism and another enriched pathway, ranked by interaction strength and FDR significance (***Fig. 4D*, *bottom)***, included retinol metabolism, oxidoreductase activity, cytokine signaling, and lipid-processing pathways, indicating coordinated metabolic and inflammatory derangements in the steatotic liver of HFpEF-MASLD mice.

Functional annotation using the DAVID bioinformatics resource further confirmed that the most significantly enriched GO terms and KEGG pathways involved retinol metabolism, fatty-acid oxidation, and inflammatory signaling cascades, which may contribute to the hepatic pathology associated with HFpEF progression in the HFpEF-MASLD model (***Fig. 4E***). Notably, several genes involved in retinoic acid synthesis and signaling were markedly downregulated, suggesting a hepatic retinoid deficiency state in HFpEF-MASLD. These findings indicate that perturbation in hepatic retinoid metabolism and accompanying inflammatory activation may constitute critical upstream events linking steatotic liver disease to cardiac diastolic dysfunction in this model.

To validate the transcriptomic findings and assess tissue-level alterations in retinoid signaling, we performed immunofluorescence staining for key proteins involved in retinol metabolism, including retinoic acid receptor α (RARα), retinoic acid receptor β (RARβ), and retinol dehydrogenase (RDH) in both liver and heart tissues from mice with HFpEF-MASLD and mice with HFpEF without MASLD. In the liver, RARα, RARβ, and RDH immunostaining (red) was localized primarily in hepatocytes, with quantitative analysis demonstrating significantly decreased expression of all three markers in the combined HFpEF-MASLD group compared to HFpEF-alone group, consistent with the transcriptomic downregulation of retinoid pathway genes (***Fig. S1A***).

Given prior studies demonstrating that retinoid signaling is disrupted in HF, including downregulation of retinol-metabolizing enzymes (34,35), we hypothesized that retinol signaling impairment in cardiac tissue may similarly contribute to HFpEF pathology in the HFpEF-MASLD model and thus elected to assess RARα, RARβ, and RDH expression in the myocardium. Interestingly, similar patterns of downregulation were observed in the heart. Retinoid-related protein staining, predominantly localized within cardiomyocytes, showed a marked reduction in the expression of RARα, RARβ, and RDH in mice with HFpEF-MASLD compared with those with HFpEF alone (***Fig. S1A***). Together, these histological data corroborate the transcriptomic results and suggest that impaired retinoid signaling may contribute to the metabolic and structural remodeling characteristic of HFpEF progression in the setting of MASLD.

### Cardiac metabolic profiling reveals perturbations in arginine biosynthesis, linoleic acid deficiency, and reduction of trigonelline in HFpEF-MASLD

To delineate myocardial metabolic alterations underlying the combined HFpEF-MASLD phenotype, targeted metabolomic profiling of left ventricular tissue was performed. A Volcano plot analysis was utilized to assess the magnitude and statistical significance of metabolite differences between the HFpEF-MASLD and HFpEF-alone groups (***Figure 5A***). This analysis identified 36 significantly altered metabolites (FDR <0.05), with a predominant shift toward downregulation in HFpEF-MASLD hearts, indicating global metabolic suppression (***Fig. 5A***). To examine the overall variation and discrimination between samples based on their metabolic profiles, multivariate modeling using sparse (sPLS-DA) and orthogonal (O-PLS-DA) partial least squares-discriminant analyses were performed. These analyses demonstrated clear separation between HFpEF-MASLD and HFpEF metabolic profiles, validating a distinct cardiac metabolomic signature (***Fig. 5B***). Permutation testing confirmed model robustness (p <0.001).

**Figure 5.**
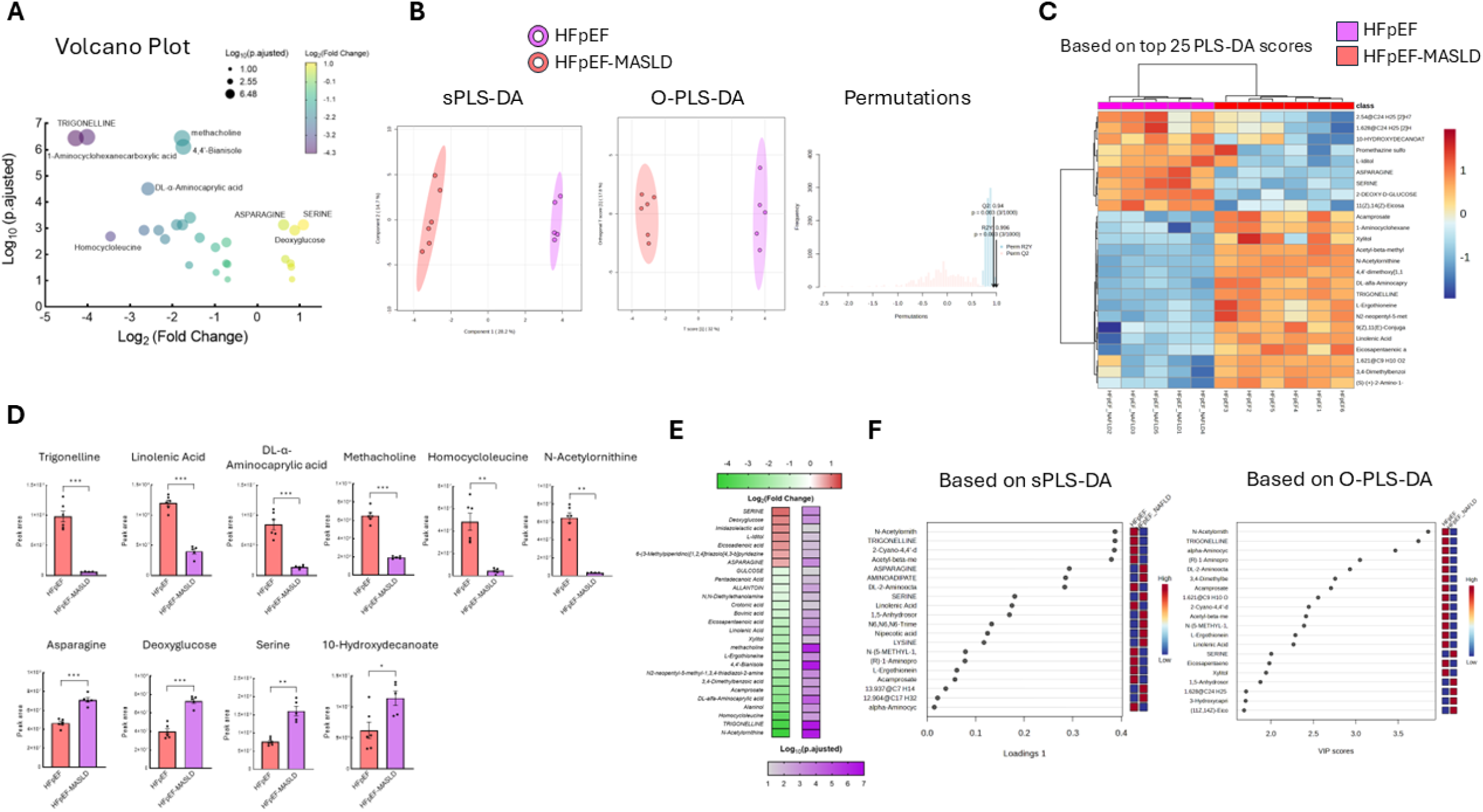
Cardiac metabolomic profiling in HFpEF-MASLD vs. HFpEF mice. **A**, Volcano plot of differentially abundant metabolites between HFpEF-MASLD and HFpEF. Points are colored by significance (FDR-adjusted p <0.05); axes show log_2_(fold change) and −log_10_(adjusted p). **B**, sPLS-DA (left) and OPLS-DA (right) score plots for HFpEF-MASLD (purple) and HFpEF (pink). Model overfitting was assessed by 1000-permutation testing (bottom); see Methods for metrics and thresholds. **C**, Heatmaps of top-25 cardiac metabolites ranked by the highest VIP (variable importance in projection) scores from PLS-DA (partial least squares discriminant analysis). Each row represents a metabolite (row z-score); each column represents a sample; clustering by Ward.D2 linkage, Euclidean distance. **D**, Relative abundance of selected significantly altered metabolites (both upregulated and downregulated). Bars represent mean ± SEM with individual values; unpaired two-tailed t-test, with Welch’s correction when variances were unequal (F-test); adjusted p-values reported where applicable. **E**, Ranked dot (lollipop) plot summarizing log_2_(fold-change) and adjusted p-value for the top differential metabolites. **F**, Loadings (Component 1) and VIP score plots from sPLS-DA/OPLS-DA for the top-20 metabolites; dashed guideline at VIP = 1. Unknown features are denoted by ‘@’.

To further visualize and identify key metabolites driving this separation, complementary hierarchical clustering heatmaps (Ward.D2 linkage; Euclidean distance) of the top 25 metabolites were generated, organizing samples based on the similarity of metabolite concentrations (***Fig. 5C***). The results clearly showed distinct clusterization of samples corresponding to the HFpEF and HFpEF-MASLD groups, further highlighting their unique metabolite profiles, as evidenced by their variable importance in projection (VIP) scores (***Fig. 5C***). Targeted inspection of the relative abundance of the most significantly altered metabolites, as analyzed by the area under the peak, revealed profound differences between the HFpEF and HFpEF-MASLD groups (***Fig. 5D***). The majority of these metabolites were decreased in the combined HFpEF-MASLD group, including trigonelline, linoleic acid, DL-α-aminocaprylic acid, methacholine, homocycloleucine, N-acetylornithine, and asparagine, whereas only a small subset of metabolites (n =4) exhibited increased levels in this group (***Fig. 5D***). To visualize both the magnitude and significance of differential metabolites, a ranked dot (lollipop) plot of log_2_(fold-change) and adjusted p values highlighted that the most substantially reduced metabolites were trigonelline, linoleic acid, N-acetylornithine, and DL-α-aminocaprylic acid (***Fig. 5E***). These findings reinforce a coordinated down-regulation of amino-acid and lipid metabolic intermediates in the HFpEF-MASLD myocardium. To determine the relative contribution of individual metabolites to the discriminant models, VIP and loading analyses were performed (***Fig. 5F***). The highest-ranking metabolites included trigonelline, linoleic acid, N-acetylornithine, and asparagine, consistent with the univariate results.

To elucidate the biological context of these altered cardiac metabolites and identify perturbed metabolic pathways, KEGG pathway enrichment analysis was performed using the MetaboAnalyst 6.0 platform. The arginine biosynthesis pathway emerged as the most significantly affected pathway, with high enrichment significance and strong topological impact (***Fig. 6A***), indicating a discernible and significant disruption in the arginine biosynthesis pathway in the myocardium of HFpEF-MASLD mice compared to HFpEF-alone mice (***Fig. 6A***). Mapping of detected metabolites onto the arginine biosynthesis pathway, which consists of 14 known metabolites, revealed coherent suppression of multiple intermediates, particularly N-acetylornithine, L-aspartate, and L-citrulline, indicating diminished flux toward L-arginine, the obligate substrate for endothelial nitric oxide (NO) bioavailability, endothelial dysfunction, and increased cardiomyocyte stiffness in the HFpEF-MASLD model (***Fig. 6B***). Collectively, these metabolomic data identify a coherent pattern of amino-acid and lipid metabolic perturbation in HFpEF-MASLD, characterized by reduced arginine biosynthesis and linoleic acid deficiency. Additionally, trigonelline, a gut microbiota-derived metabolite, was among the significantly decreased metabolites in HFpEF-MASLD hearts, indicating possible alterations in gut-derived metabolic pathways in this model.

**Figure 6:**
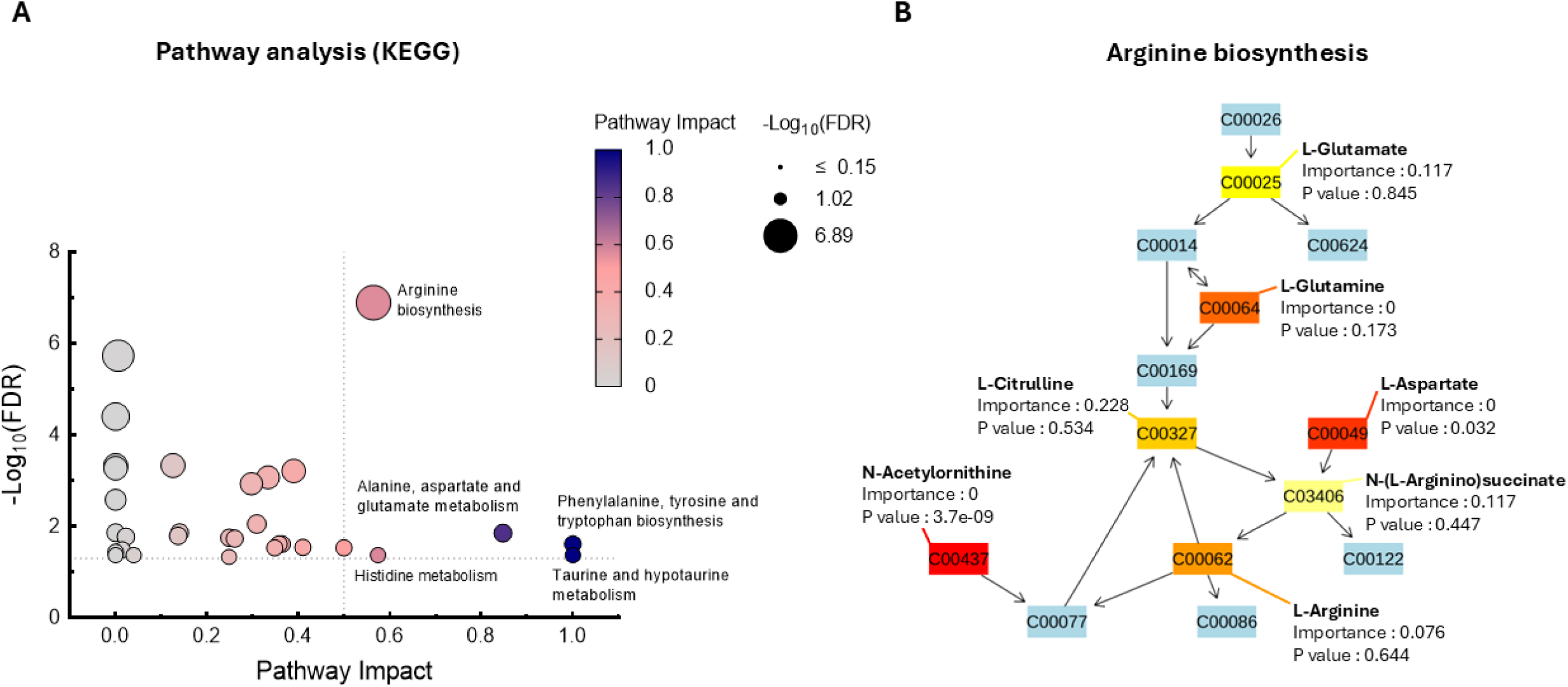
Metabolic pathway analysis of cardiac metabolites distinguishing HFpEF-MASLD from HFpEF. **A,** KEGG pathway enrichment bubble plot (MetaboAnalyst 6.0; species =*Mus musculus*). X-axis: pathway impact (topology, relative-betweenness centrality). Y-axis: −log_10_(FDR). Bubble size is proportional to the number of metabolite hits in the pathway; color encodes −log_10_(FDR) (darker = more significant). Enrichment by hypergeometric test with Benjamini-Hochberg correction. **B,** Pathway topology diagram for arginine metabolism. Node fill indicates mean log_2_ fold-change (HFpEF-MASLD vs. HFpEF); blue nodes denote KEGG metabolites not detected in our dataset. The Importance value reflects node centrality (relative-betweenness) within the pathway; node-level *p* values derive from the pathway analysis. Directed edges correspond to KEGG reactions.

## Discussion

The present study provides experimental evidence that MASLD directly exacerbates HFpEF pathophysiology. Even in the absence of overt diabetes or hypertension, MASLD markedly worsened diastolic function and cardiac remodeling in our pressure-overload HFpEF model. HFpEF-MASLD mice exhibited pronounced reductions in diastolic relaxation (lower septal e′ velocity) and higher filling pressures (elevated E/e′ ratio) despite preserved ejection fraction, along with accentuated ventricular hypertrophy and fibrosis compared with HFpEF alone. This study suggests that while the full spectrum of metabolic syndrome did not manifest, the presence of MASLD was still associated with measurable impairment of cardiac diastolic properties. These findings align with clinical evidence demonstrating that MASLD can independently impair cardiac diastolic function, with elevated left ventricular filling pressures and relaxation abnormalities even in non-obese or metabolically “normal” patients (19,20, 22). Large-scale epidemiological studies further associate MASLD with higher risk of incident HF and worse prognosis, predominantly HFpEF (7,8,36–38). Indeed, the majority of MASLD patients who develop HF present with preserved ejection fraction (8,38). Collectively, these data underscore the clinical significance of the liver-heart axis and support a causal role for MASLD in promoting HFpEF progression.

Mechanistically, our study implicates two principal pathways by which MASLD may accelerate HFpEF: chronic inflammation and retinoid metabolic dysregulation. First, we observed robust myocardial immune infiltration in HFpEF-MASLD mice, with significant increases in CD3⁺ T cells and CD68⁺ macrophages compared with HFpEF alone. This finding highlights the proinflammatory state imposed by MASLD, consistent with the established role of chronic low-grade inflammation in HFpEF progression (12,13,39–41). Cardiac immune activation has been linked to mitochondrial dysfunction, impaired oxidative phosphorylation, and maladaptive fibrosis (12,39–41). Similarly, obesity-associated macrophage activation through the NLRP3 inflammasome promotes sympathetic overactivity and cardiac remodeling (39). These findings, together with our data, suggest that MASLD amplifies myocardial inflammation and metabolic stress through systemic immune activation, a central mechanism underlying the “metabolic HFpEF” phenotype (12,13,39–41). This cardio-hepatic inflammatory axis may therefore represent a key driver of HFpEF severity and progression in the context of steatotic liver disease.

The second major mechanistic insight from our study is the disruption of retinol (vitamin A) metabolism as a potential molecular link between hepatic steatosis and cardiac dysfunction. Transcriptomic and histological analyses demonstrated downregulation of retinoic acid signaling in both the liver and heart of HFpEF-MASLD mice, with decreased expression of RARα, RARβ, and RDH enzymes. Retinoic acid serves as a critical regulator of gene transcription in myocardial development and homeostasis, and its deficiency has been associated with cardiac hypertrophy, fibrosis, and diastolic dysfunction (34,42–47). Recent evidence indicates that failing human hearts exhibit up to a 40% reduction in myocardial all-trans retinoic acid levels (34). Experimental disruption of retinoid signaling, such as RARα knockout, leads to delayed myocardial relaxation and elevated filling pressures, recapitulating an HFpEF-like phenotype (34,42–44). Conversely, pharmacologic activation of retinoic acid signaling has been shown to attenuate cardiac fibrosis and improve diastolic performance in preclinical models (35,45,46). Our data extend these findings by suggesting that hepatic retinoid dysregulation in MASLD can propagate to the heart, creating a systemic retinoid deficiency that contributes to diastolic dysfunction.

Metabolomic profiling further revealed a downregulation of arginine biosynthesis in the hearts of HFpEF-MASLD mice. Arginine (L-arginine) serves as the obligate substrate for endothelial nitric oxide synthase (eNOS) to generate nitric oxide (NO), and reduced arginine availability constrains NO synthesis, thereby undermining endothelial vasodilatory and anti-inflammatory functions (47,48). In HFpEF, coronary microvascular endothelial dysfunction is a hallmark pathophysiological feature that limits NO bioavailability and impairs NO-cyclic guanine monophosphate (cGMP)-protein kinase G (PKG) signaling cascade (49–52). Attenuation of this pathway diminishes titin phosphorylation, leading to increased cardiomyocyte stiffness and diastolic dysfunction (51,52). Both preclinical and translational HFpEF studies have demonstrated dysregulation of arginine metabolism, including increased arginase activity and accumulation of asymmetric dimethylarginine (ADMA), which competes for or inhibits eNOS, as key drivers of reduced NO bioavailability and endothelial dysfunction (48,50,53). These findings establish a mechanistic link between arginine deficiency, endothelial dysfunction, and myocardial stiffening in HFpEF. Beyond its vascular role, arginine also functions as an auxiliary energy substrate under hypoxic conditions, providing metabolic flexibility during cardiac stress. Moreover, loss of retinoid signaling has been shown to repress enzymes involved in arginine biosynthesis, resulting in decreased myocardial arginine content (34,35). Together, these observations support a novel vitamin A-NO metabolic axis in MASLD-driven HFpEF, whereby hepatic retinoid impairment may propagate systemic NO insufficiency and diastolic dysfunction.

The observed reduction in myocardial linoleic acid in HFpEF-MASLD mice may represent another important metabolic link between hepatic steatosis and mitochondrial dysfunction in the heart. Linoleic acid is the major polyunsaturated fatty acid moiety of cardiolipin, a mitochondrial inner-membrane phospholipid essential for electron-transport chain supercomplex assembly and mitochondrial oxidative phosphorylation. Thus, linoleic acid is essential for maintaining mitochondrial integrity and bioenergetic efficiency. Indeed, depletion of linoleic acid has been shown to reduce cardiolipin content, destabilize electron-transport chain complexes, and impair mitochondrial oxidative phosphorylation; mechanisms consistently implicated in the pathophysiology of HF (54,55). Experimental evidence has demonstrated that dietary linoleate supplementation preserves cardiolipin composition and attenuates mitochondrial dysfunction in failing hearts (56). Moreover, linoleic acid promotes proper assembly of mitochondrial respiratory complexes II, III_2_ and IV, thereby improving electron transport and ATP generation (57). In the HFpEF-MASLD model, the marked decrease in linoleic acid may therefore exacerbate myocardial bioenergetic insufficiency, diminish mitochondrial reserve capacity, and impair relaxation. Given the systemic metabolic stress imposed by MASLD, linoleate depletion may further propagate mitochondrial injury and oxidative stress, contributing to HFpEF progression through a cardio-hepatic metabolic axis.

In addition, our metabolomic analysis identified a marked reduction in trigonelline levels in HFpEF-MASLD mice. Trigonelline is a bioactive alkaloid derived from dietary sources and gut microbial metabolism, known for its anti-inflammatory, antioxidative, and vasoprotective properties (58–60). Mechanistically, trigonelline inhibits the intestinal conversion of choline to trimethylamine (TMA) and its subsequent hepatic oxidation to trimethylamine-N-oxide (TMAO), a gut-derived metabolite strongly linked to atherosclerosis, HF risk, and poor HF prognosis (58, 61–63). Experimental models have demonstrated that dietary choline or TMAO supplementation aggravates pressure-overload-induced cardiac dysfunction and myocardial fibrosis, highlighting a direct role of the TMAO pathway in HFpEF progression (64). Beyond its effects on lipid metabolism, trigonelline has been shown to enhance endothelial NO production and improve vascular reactivity (60). Thus, the pronounced decline in trigonelline levels observed in our HFpEF-MASLD model may exacerbate endothelial dysfunction and diastolic impairment, further contributing to the pathogenesis of this combined phenotype. These findings suggest that perturbations in gut-liver-heart metabolic signaling may represent an additional mechanistic layer in MASLD-associated HFpEF pathogenesis.

Collectively, our results indicate that MASLD worsens HFpEF through synergistic metabolic and inflammatory mechanisms. The proposed model illustrates how steatotic liver disease promotes systemic inflammation and perturbs retinoid and arginine/NO pathways, leading to enhanced myocardial fibrosis, hypertrophy, and impaired relaxation. These insights reinforce the need for early recognition of HFpEF risk in the setting of MASLD and vice versa. Identification of this “metabolic-inflammatory” HFpEF phenotype could guide tailored interventions aimed at both hepatic and cardiac targets. Therapeutic strategies directed at restoring retinoid signaling, augmenting NO bioavailability, or modulating gut-derived metabolites such as trigonelline warrant further exploration. Given that nearly half of HFpEF patients exhibit features of metabolic liver disease (10,15,24), interventions targeting the underlying hepatic dysfunction, through weight loss, anti-inflammatory therapies, or metabolic agents such as GLP-1 receptor agonists and SGLT2 inhibitors, may offer dual benefits on both liver and heart outcomes (65). Future translational studies should assess cardiac retinoid signaling, NO pathway integrity, and metabolomic profiles in HFpEF patients with MASLD to validate these mechanisms and identify biomarkers for risk stratification and therapy.

This study has several limitations. First, it was conducted in a murine model combining pressure overload and diet-induced MASLD, which, although recapitulating hallmark HFpEF features, may not fully represent the human disease spectrum. Second, our MASLD model reflects an early stage of hepatic steatosis without advanced fibrosis; therefore, the cardio-hepatic interactions in later stages may differ, and this model may not fully capture the human HFpEF-MASLD continuum. Third, metabolic and transcriptomic analyses were cross-sectional and hypothesis-generating; causal inference was not tested through pathway-specific interventions, such as retinoid or arginine supplementation. Finally, validation in human HFpEF-MASLD cohorts is required to confirm transitional relevance. Despite these limitations, the integrated phenotyping, transcriptomic, and metabolomic data provide mechanistic insights into MASLD amplifies HFpEF pathophysiology.

In conclusion, MASLD amplifies the severity of HFpEF by integrating inflammatory and metabolic stress across the liver-heart axis. Our findings identify retinoid pathway dysregulation and arginine-NO metabolism as central mechanistic links, providing a rationale for targeted, mechanism-based interventions in metabolic HFpEF.

## Abbreviations

ADMA: asymmetric dimethylarginine;
ANOVA: analysis of variance;
ATRA: all-trans retinoic acid;
cGMP: cyclic guanosine monophosphate;
DAPI: 4′,6-diamidino-2-phenylindole;
EF: ejection fraction;
GLS: global longitudinal strain;
HF: heart failure;
HFpEF: heart failure with preserved ejection fraction;
H&E: hematoxylin and eosin;
HFD: high-fat diet;
KEGG: Kyoto Encyclopedia of Genes and Genomes;
LV: left ventricle/ventricular;
LVEDD: left-ventricular end-diastolic diameter;
LVESD: left-ventricular end-systolic diameter;
MASLD: metabolic dysfunction-associated steatotic liver disease;
MASH: metabolic dysfunction-associated steatohepatitis;
NO: nitric oxide;
NOS: nitric oxide synthase;
PKG: protein kinase G;
PLS-DA: partial least-squares discriminant analysis;
RAR: retinoic acid receptor;
RDH: retinol dehydrogenase;
RNA-seq: RNA sequencing;
SEM: standard error of the mean;
TAC: transverse aortic constriction;
TMA: trimethylamine;
TMAO: trimethylamine-N-oxide;
UHPLC-MS/MS: ultra-high-performance liquid chromatography-tandem mass spectrometry.

## Acknowledgments

Funding Support and Author Disclosures

The authors report no relationships relevant to the contents of this paper to disclose.

---

**Figure S1.**
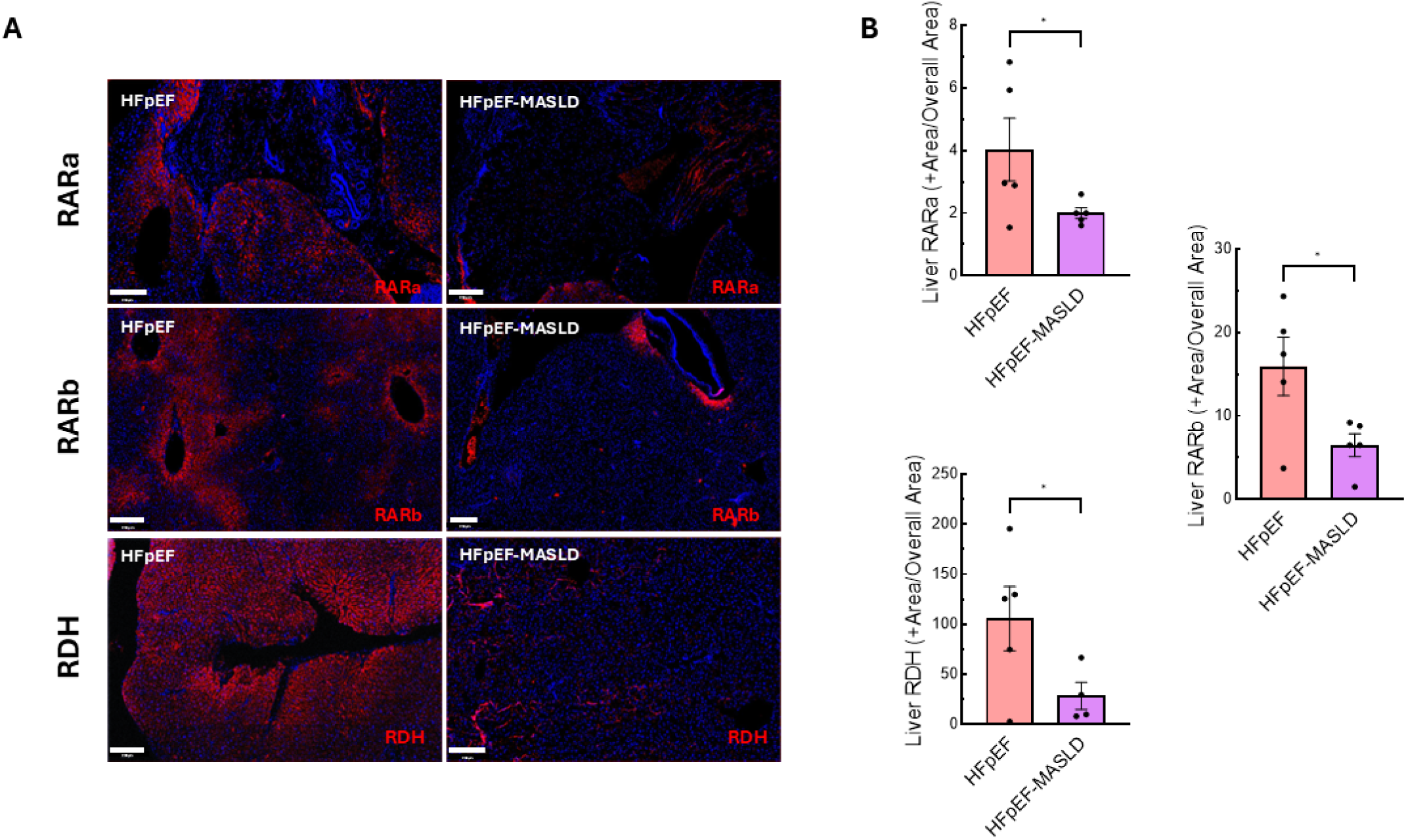
Immunofluorescent staining of hepatic retinoid-signaling proteins in HFpEF and HFpEF-MASLD mice. **A**, Representative liver sections stained for retinoic-acid receptor α (RARα, top), retinoic-acid receptor β (RARβ, middle), and retinol dehydrogenase (RDH, bottom). Red = specific marker; nuclei = DAPI (blue). Scale bar = 100 µm; magnification ×20. **B**, Quantitative analysis of RARα⁺, RARβ⁺, and RDH⁺ immunoreactive cells per tissue area. Data are mean ± SEM; individual mice shown as dots (n = 6 per group).

**Figure S2.**
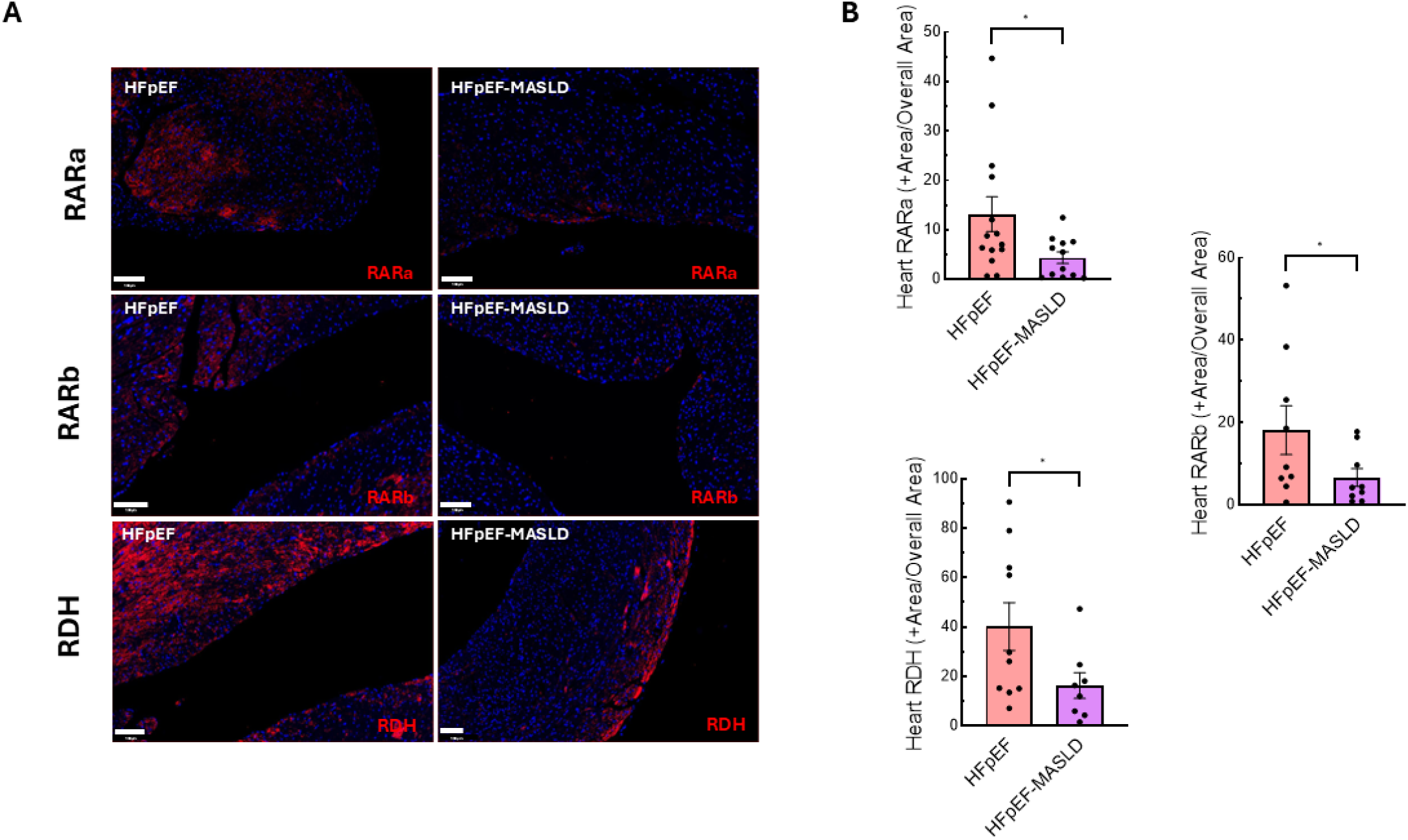
Immunofluorescent staining of myocardial retinoid-signaling proteins in HFpEF and HFpEF-MASLD mice. **A,** Representative left-ventricular sections stained for RARα (top), RARβ (middle), and RDH (bottom). Red = specific marker; nuclei = DAPI (blue). Scale bar = 100 µm; magnification ×20. **B**, Quantification of RARα⁺, RARβ⁺, and RDH⁺ cardiomyocytes per tissue area. Data are mean ± SEM; individual mice shown as dots (n = 6 per group).

